# Disrupting the Right Inferior Frontal Gyrus Increases ‘Social Brain’ Activity During Emotion Recognition

**DOI:** 10.64898/2026.10.08.757563

**Authors:** Emel Küçük, Hannah Gofton, Cameron Smith, Daniel Kaiser, David Pitcher

**Affiliations:** Department of Psychology, University of York, Heslington, York, YO10 5DD, U.K; Neural Computation Group, Department of Mathematics and Computer Science, Physics, Geography, Justus Liebig University Giessen, 35392 Giessen, Germany; Center for Applied Computer Science and Data Science (ZAD), Justus Liebig University Giessen, 35392 Giessen, Germany; Center for Mind, Brain and Behavior (CMBB), Universities of Marburg, Giessen, and Darmstadt, 35032 Marburg, Germany; Cluster of Excellence “The Adaptive Mind”, Universities of Giessen, Marburg, and Darmstadt, 35394 Giessen, Germany

**Keywords:** Thetaburst transcranial magnetic stimulation (TBS), functional magnetic resonance imaging (fMRI), emotion recognition

## Abstract

Neuroimaging studies have identified a specialised social brain network that supports emotion recognition and social cognition. This network performs a range of functions that include facial expression recognition and inferring the mental states of others. Studies of neuropsychological patients report that damage to the right inferior frontal gyrus (IFG), the top-down control area in this network, impairs facial expression recognition. To investigate how transient disruption of the right IFG affects the wider social brain we combined transcranial magnetic stimulation (TMS) with functional magnetic resonance imaging (fMRI). Participants (*n*=20) were scanned across two sessions while performing a delayed match-to-sample emotion recognition task in which happy or fearful expressions were preceded by a congruent or incongruent word cue. Prior to each scan, thetaburst TMS was delivered over the right IFG or over the vertex control site (session order was balanced across participants). A group whole-brain analysis revealed that TMS to the IFG increased neural activity in multiple regions of the social brain network. These included the bilateral temporal pole, precuneus and medial prefrontal cortex (mPFC). This pattern of results shows that the disruption of top-down control mechanisms localised in the IFG alters the neural processing in remote nodes of the social brain network. Our findings are discussed within the wider context of how we recognise and understand the emotions of other people.

## 1 Introduction

Neuroimaging studies have identified a distributed network of brain regions that work together to support the social interactions that are the basis of our daily lives (Amodio & Frith, 2006; Brothers, 1990; Frith & Frith, 2007). A core function of the social brain network is to predict the actions of other people based on our perception and understanding of their mental and emotional state. The more successful the prediction, the more successful the subsequent social interaction. Such a complex cognitive task requires input from a network of brain regions with different functional roles. The medial prefrontal cortex (mPFC) and the temporo-parietal junction (TPJ) represent the mental states of others (Saxe & Kanwisher, 2013), what is also called theory of mind (Premack & Woodruff, 1978). The posterior superior temporal sulcus (pSTS) and the amygdala mediate the recognition and categorisation of emotions based on facial expressions (Morris et al., 1996; Pitcher, 2014; Sliwinska & Pitcher, 2018). The temporal poles store biographical and semantic information that is used to contextualise behaviour (Spreng & Levine, 2006). How does this network of regions coordinate to organize and interpret perceptual information for accurately inferring the emotional states of others? It has been suggested that the lateral prefrontal cortex (PFC) exerts top-down control over face-selective perceptual areas to facilitate adaptive emotion recognition (Pitcher et al., 2023; Wang et al., 2020). In the current study, we tested how a transient disruption of the right lateral PFC impacts the neural correlates of face emotion recognition. To this end, we combined transcranial magnetic stimulation (TMS) with functional magnetic resonance imaging (fMRI).

The lateral PFC is implicated in several neural processes that support our understanding of other people. These include cognitive control (MacDonald et al., 2000), working memory (Curtis & D’Esposito, 2003), theory of mind (Kalbe et al., 2010; Rowe et al., 2001) and executive function (Goldman-Rakic, 1996). Importantly a part of the lateral PFC called the inferior frontal gyrus (IFG) has been associated with many aspects of social cognition, including emotion perception (Kilts et al., 2003), social judgements (Bzdok et al., 2012) and understanding the intentions of others (Liu et al., 2016). The face is a salient cue that underpins all these functions and studies show that the IFG demonstrates selectivity towards faces and socially relevant cues (Chan & Downing, 2011; Davies-Thompson & Andrews, 2012; Nikel et al., 2022; Tso et al., 2018). Furthermore, studies have shown that the IFG exhibits a modulatory role in emotion recognition through highlighting a task-dependent top-down influence over perceptual brain regions. For example, by directing selective attention towards the identity or emotion of a face, the IFG can modulate the response of the FFA and STS, respectively (Baldauf & Desimone, 2014; Bzdok et al., 2012). This top-down control allows the IFG to flexibly modulate stimulus-driven, bottom-up processing, supporting the efficient and adaptive processing of social cues.

The broader involvement of the IFG in a wide range of processes including language (Calvert & Campbell, 2003; Calvert et al., 2000; Romanski, 2012) and cognitive control (Aron et al., 2004; Wager & Smith, 2003) builds upon its anatomical and functional connections with other nodes of the social brain. Direct anatomical connections have been found between the STS and IFG via the arcuate fasciculus (AF), a white matter tract connecting frontal, temporal and parietal brain regions (Rilling et al., 2008). Disruptions to the AF have been related to deficits in emotion recognition and social understanding (Herbet et al., 2014; Nakajima et al., 2018). Moreover, using a developmental approach by comparing children with and without false belief understanding, Grosse Wiesmann et al. (2017) found a significant correlation between the structural integrity of the AF and Theory of Mind (ToM) performance. These findings suggest that the emergence of ToM abilities during childhood are supported by the extent to which temporoparietal regions are connected to the IFG via the AF. Similarly, Parkinson and Wheatley (2014) found a positive correlation between emotional empathy and white matter integrity of the AF. This suggests that more robust connections between areas involved in emotion understanding and emotion perception are associated with greater levels of emotional empathy. This pattern of results is also consistent with the hypothesis that the STS and IFG are components in a functional network specialised for processing dynamic aspects of social perception (Pitcher, 2025; Pitcher & Ungerleider, 2021; Wang et al., 2020).

Brain lesion studies have found that damage to the lateral prefrontal cortex, particularly to the IFG, is associated with reduced accuracy in emotion recognition tasks (Adolphs et al., 2000; Philippi et al., 2009). Adolphs et al. (2000) reported an association between lesion density and task performance in patients with focal lesions in the frontal cortex (including the IFG) when recognising and naming emotional facial expressions. Similarly, Philippi et al. (2009) showed patients faces depicting six basic emotions and found that lesions involving the IFG were associated with deficits in emotion recognition. Neuropsychological evidence from studies using the Reading the Mind in the Eyes Test (RMET) further support the role of the IFG in emotion recognition. In the RMET, photographs of eye regions are presented, and participants indicate which of four adjectives best describes the mental state depicted in the image. When completing this task, patients with lesions in the IFG perform significantly worse than patients with lesions excluding the IFG and healthy controls (Dal Monte et al., 2014; Perry et al., 2017). Consistent with the patient data, TMS delivered over the right dorsolateral prefrontal cortex (dlPFC) resulted in slower reaction times for fearful but not neutral faces in a facial expression identification task, whereas there was no effect of TMS on a gender discrimination control task (Notzon et al., 2018; Zwanzger et al., 2014). More recently we demonstrated that the thetaburst (TBS) TMS delivered over the right PFC increased activity in regions of the Default Mode Network (DMN) while participants performed a facial expression naming task (Pitcher et al., 2023). This result demonstrated brain networks with different functions can exhibit compensatory neural activity following transient disruption of the PFC.

In the current study we combined TMS and fMRI to investigate how disrupting the prefrontal cortex impacts neural activity in other brain regions when recognising fearful and happy facial expressions. Across two sessions (completed on different days) participants performed a delayed match-to-sample emotion recognition task where the word happy or fearful was presented, followed by an Ekman face displaying a happy or fearful facial expression (Ekman & Friesen, 1976). Thetaburst (TBS) TMS was delivered over the face-selective area in the right IFG (Nikel et al., 2022) or the vertex (session order was balanced across participants).

## 2 Methods

### 2.1 Participants

Twenty participants (14 females; age range 19-42 years; mean age 22.40, SD = 4.91) gave written informed consent as directed by the Ethics committee at the University of York. All participants had normal or corrected-to-normal vision and no history of neurological disorders.

### 2.2 Materials

#### 2.2.1 Experimental Task

Stimuli consisted of photographs from the Pictures of Facial Affect collection (Ekman & Friesen, 1976). From this collection, a total of 22 faces were selected. These faces included 11 identities (6 female, 5 male) each depicting a happy and fearful expression. The words ‘Happy’ or ‘Fearful’ were presented in black text. All stimuli were presented on a grey background in the centre of the screen.

#### 2.2.2 Functional Localiser

To identify face-selective brain regions, participants viewed clips of faces and objects that have been used in prior studies (Gonzalez Alam et al., 2025; Küçük et al., 2024; Sliwinska et al., 2022). This functional localiser included 60 3-second movie clips for each of the following categories: faces, bodies, objects, scenes, scrambled objects. The face and body clips were filmed against a black background, capturing only the faces or bodies of seven different children as they individually danced or played (with toys or adults that were out of frame). Object stimuli consisted of 15 different moving objects that minimised any impression of animacy or external manipulation (e.g. balls rolling down slopes). Scene stimuli were filmed primarily from a car window moving across 15 different locations (e.g. pastoral scenes).

Scrambled objects were generated by dividing each object movie clip into a 15x15 grid and spatially rearranging each frame. Rest blocks were also included, during which the screen displayed a randomly selected colour that changed every 3 seconds. All stimuli were presented in full screen at a resolution of 1920x1080 pixels.

### 2.3 Design and Procedure

Participants completed two separate TMS sessions that were performed on different days (ranging from 1-19 days apart; averaging 7.2 days; SD = 4.74). Offline TBS was delivered over the functionally localised face area in the right IFG in one session and over the vertex (control site) in the other session. The order of stimulation sites was balanced across participants. Before the fMRI scan, the Brainsight TMS-MRI co-registration system (Rogue Research) was used to mark the location of the face-selective area in the right IFG based on each participant’s fMRI localiser data. The vertex control site was identified, using a tape measure, as the midpoint between the nasion and inion. Participants were then taken to the fMRI scanner control room where thetaburst TMS was delivered over the right IFG or vertex. Since the effects of TMS are transient, participants entered the scanner and functional data collection began within five minutes of TMS administration. After the final session, participants were debriefed on the nature of the study.

#### 2.3.1 Experimental Task

Functional data for the delayed match-to-sample emotion recognition task were acquired across three runs, each lasting 460 seconds, including an additional 10 seconds at the start and end of the run. Each run consisted of 88 randomised trials (illustrated in Figure 1). Trials began with a fixation cross presented in the centre of the screen for 500ms, followed by a 1-second presentation of an affective label (“happy” or “fearful”), a blank screen for 500ms, then a 3-second display of a face depicting either a happy or fearful expression. Participants were instructed to indicate whether the word ‘happy’ or ‘fearful’ matched the facial expression that followed. In each experimental run, there were 22 trials for each of the four conditions: fearful congruent, fearful incongruent, happy congruent and happy incongruent. All experimental data were collected within 30 minutes of TMS being delivered. Prior studies have demonstrated that this time window is effective for capturing TMS-induced changes on the BOLD signal (Groen et al., 2021; Handwerker et al., 2020; Pitcher et al., 2014; Pitcher et al., 2017).

**Figure 1.**
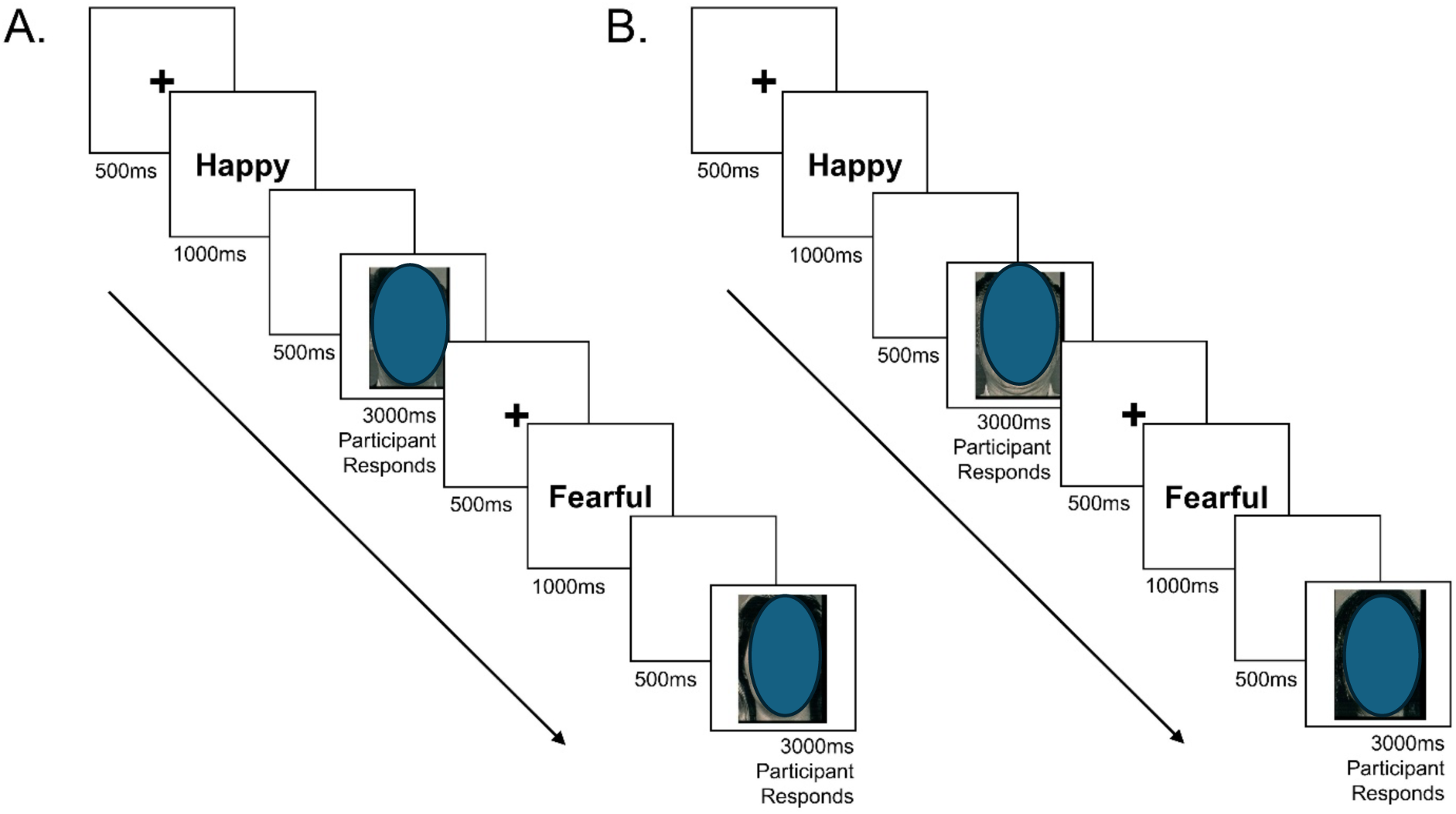
The Trial Procedure for the Delayed Match-to-Sample Task. The examples here demonstrate both congruent (A) and incongruent trials (B) from the delayed match-to-sample task which was completed in the MRI scanner. Participants indicated whether the facial expression matched the previously presented affective word, and their response was taken whilst the face was displayed on the screen.

#### 2.3.2 Functional Localiser

Upon completing the experimental task, participants completed two runs of the localiser task to individually identify face-selective areas. Data were acquired over blocked-design functional runs, each lasting 234 seconds, including an additional 10 seconds at the start and end of the run. Each functional run contained 2 sets of 5 consecutive stimulus blocks (faces, bodies, scenes, objects, or scrambled objects) and three rest blocks, at the beginning, middle, and end of the run. Each block lasted 18 seconds, containing six 3-second video clips from that category. The order of the stimulus category blocks in each run followed a palindromic pattern of rest, faces, bodies, scenes, objects, scrambled objects, rest, scrambled objects, objects, scenes, bodies, faces, rest. In each block, the stimuli were randomly selected from that specific category. Participants were instructed to watch the video clips but were not asked to perform any explicit task.

### 2.4 Data Acquisition

#### 2.4.1 TMS Procedure

For each participant, stimulation sites were localised using their individual structural and functional images obtained during the fMRI functional localiser task containing dynamic stimuli used in previous studies of the face network (Pitcher et al., 2011; Sliwinska et al., 2020; Sliwinska et al., 2026). The targeted stimulation site in the right IFG (Nikel et al., 2022) was identified as the peak voxel in the face-selective ROI using a contrast showing greater activation to dynamic faces compared to dynamic objects (mean MNI coordinates: 43, 11, 37). The control site (vertex) was identified as the midpoint on the top of the head, between the tip of the nose (nasion) and back of the head the head (inion). The Brainsight TMS-MRI co-registration system (Rogue Research) was used to identify the TMS sites, and the positions of the sites were marked on each participant’s scalp using a marker pen.

Using a Magstim Super Rapid Stimulator (Magstim; Whitland, UK) with a figure-eight coil (70mm wing diameter), TMS was delivered at a fixed intensity of 45% of the maximum stimulator output over the functionally localized right IFG or vertex. Continuous thetaburst stimulation was administered using a train of 600 pulses delivered as 200 bursts of 3 pulses at 30Hz with a burst frequency of 6Hz, lasting 33.3 seconds in total. We used a modified thetaburst protocol (Nyffeler et al., 2006; Sliwinska et al., 2020; Sliwinska et al., 2021) which has been shown to produce longer lasting effects than the standard protocol (Goldsworthy et al., 2012; Huang et al., 2005). During stimulation, the coil handle was held pointing upwards and parallel to the midline when delivered over the right IFG and flat against the scalp with the handle pointing towards the inion when delivered over the vertex.

#### 2.4.2 fMRI Procedure

Imaging data were acquired using a 3T Siemens Magnetom Prisma MRI scanner (Siemens Healthcare, Erlangen, Germany) with a twenty-channel phased array head coil at the University of York. Functional images were acquired with a gradient-echo EPI sequence with the following acquisition parameters providing whole brain coverage: Matrix size = 80x80, field of view (FOV) 240mm, 60 slices with 3mm slice thickness, repetition time (TR) = 2000ms, echo time (TE) = 30ms, flip angle = 80 degrees; voxel size 3mm isotropic. Acquisition order was interleaved, acquiring 230 volumes per run and slices were aligned with the anterior to posterior commissure line.

Structural images were acquired using a high-resolution T-1 weighted Magnetization Prepared Rapid-Gradient Echo (MPRAGE) sequence with the following parameters: Matrix size = 256x256, FOV 256mm, 176 slices with 1mm slice thickness, TR = 2300ms, TE = 2.26ms, inversion time (TI) = 900ms, flip angle = 8 degrees; voxel size 1mm isotropic. Acquisition order was ascending, and slices were aligned with the anterior to posterior commissure line.

### 2.5 Data Analysis

Functional MRI data were analysed using Analysis of Functional NeuroImages (AFNI; http://afni.nimh.nih.gov/afni) (Cox, 1996). The functional images were slice-time corrected and aligned to a volume with minimal outliers, then co-registered to the anatomical image. Spatial smoothing was applied to the functional data using a 5mm full-width at half-maximum (FWHM) Gaussian kernel and time points with motion exceeding 0.3mm were censored. The signal intensity for each voxel was scaled to a mean of 100.

For the functional localiser task, the GLM included five regressors of interest (faces, bodies, scenes, objects and scrambled objects) and motion parameters were included as regressors of no interest. Face-selective areas were identified in each participant using a contrast of greater activation for faces compared to objects. For the experimental task, a general linear model (GLM) was established by convolving the standard hemodynamic response function with the four experimental conditions as regressors of interest (fearful congruent, fearful incongruent, happy congruent and happy incongruent). Regressors of no interest such as motion parameters were also included to account for residual movement-related noise.

To examine the whole-brain effects of TMS on emotion recognition, a 3-way mixed-effects ANOVA with two TMS conditions (stimulation of the right IFG vs. vertex), four stimulus conditions (fearful congruent, fearful incongruent, happy congruent, happy incongruent) and twenty participants was conducted at the group level using AFNI’s 3dANOVA3. Each participant contributed statistical maps for each condition from both TMS sessions. Group whole-brain maps were calculated for each TMS session and data from the right IFG session was subtracted from the vertex session. Using 3dcalc, a group brain mask was applied to the resulting statistical dataset to exclude non-brain voxels and produce a clean dataset.

## 3 Results

The functional activation maps were thresholded at a voxel-wise uncorrected p-value of *p* < 0.005 (*z* = 3.1) with a cluster extent threshold of 40 contiguous voxels. These thresholded maps were overlaid on the MNI152 template for anatomical reference. The results revealed four regions showing significantly greater activation following TMS over the right IFG compared to the vertex (see Figure 2). Specifically, increased activation was found in the medial prefrontal cortex (mPFC; 304 voxels; peak MNI co-ordinates at -2, 64, 25), precuneus (202 voxels; peak MNI co-ordinates at 1, -53, 22), the left anterior STS (165 voxels; peak MNI co-ordinates at -68, -17, -5), and the right anterior STS (132 voxels; peak MNI co-ordinates at 64, -2, -11).

**Figure 2.**
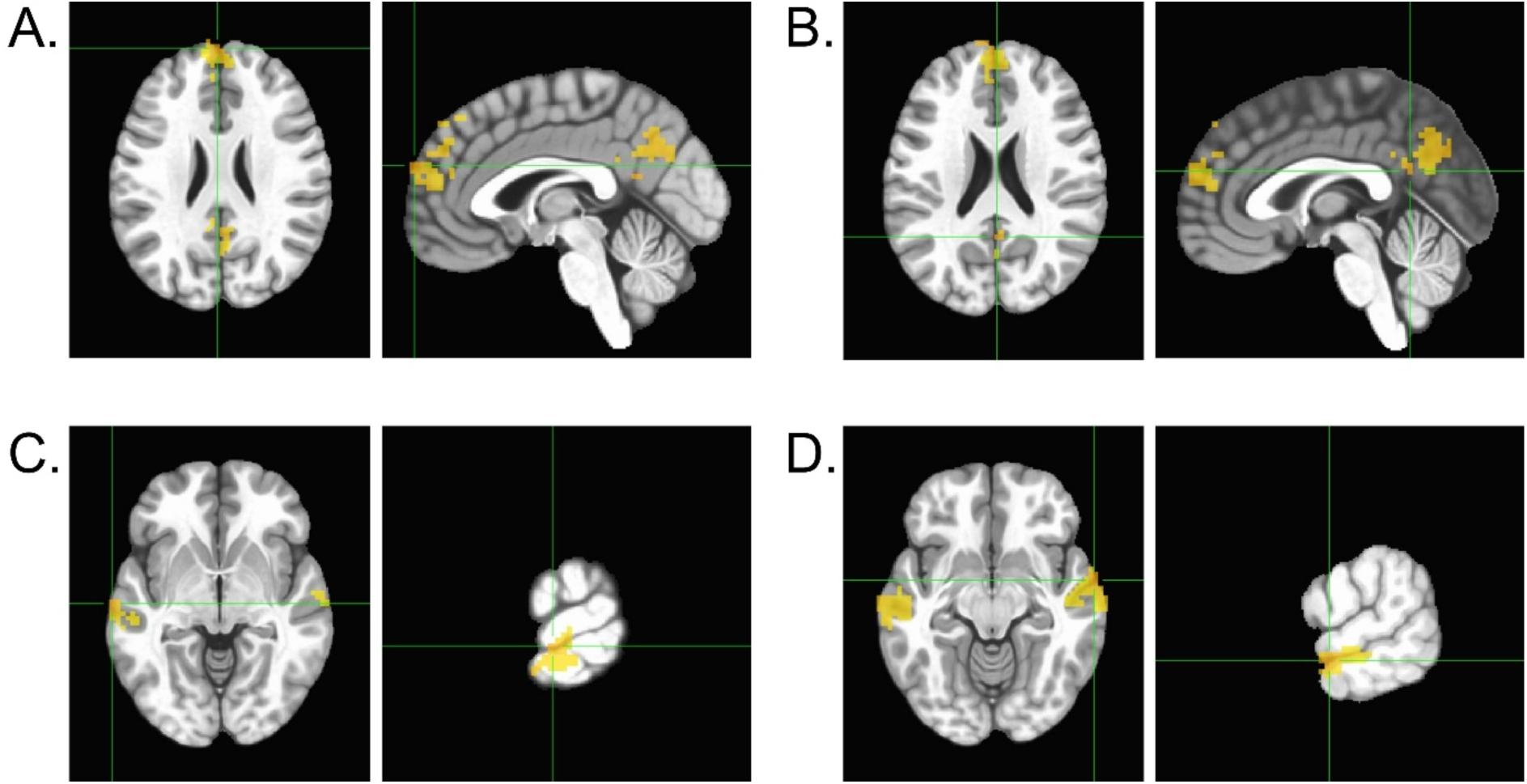
Brain Areas Showing Greater Activation When Transcranial Magnetic Stimulation Was Delivered to the Right Inferior Frontal Gyrus. Axial and sagittal slices showing clusters of significantly increased BOLD response when TMS was delivered to the right IFG compared to the vertex in the (A) medial prefrontal cortex (mPFC), (B) precuneus, (C) left anterior superior temporal sulcus (aSTS) and (D) right aSTS. These data were acquired during a delayed match-to-sample emotion recognition task. The green crosshairs mark the peak voxel within each cluster.

To determine condition-specific response magnitudes, we extracted the mean percent signal change estimates for each trial condition in each TMS session for the four key regions identified in the whole-brain analyses (see Figure 3).

**Figure 3.**
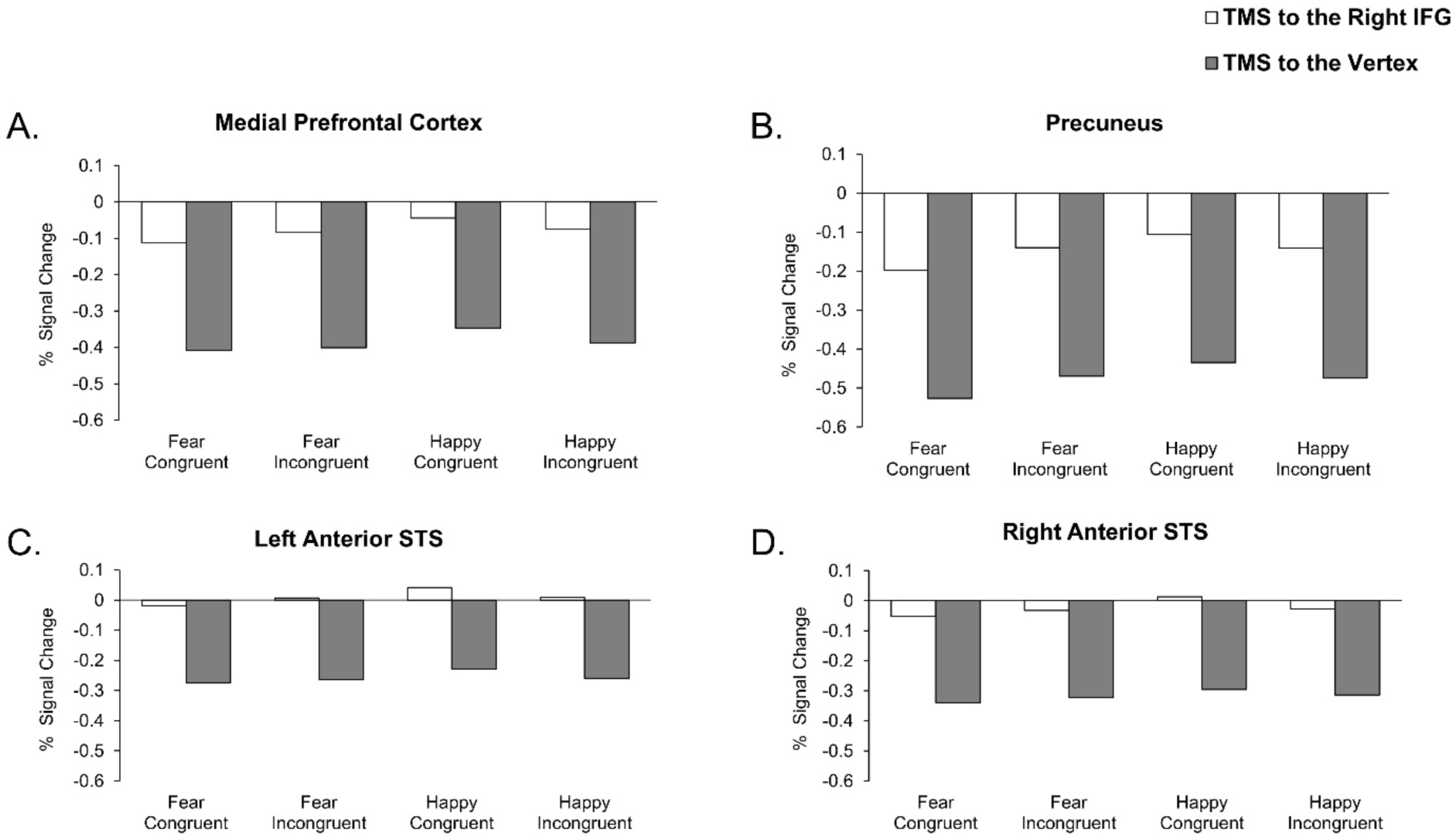
Percent Signal Change for Each Trial Condition Following TMS to the Right Inferior Frontal Gyrus and the Vertex. Bar graphs display the mean percent signal change in the mPFC, precuneus, left anterior superior temporal sulcus (aSTS) and (D) right aSTS for each trial condition (Fear Congruent, Fear Incongruent, Happy Congruent, Happy Incongruent) following TMS to the right IFG and vertex. Data are presented to illustrate activation patterns across conditions and TMS sites. TMS delivered over the right IFG resulted in increased BOLD responses in all four regions identified in the whole-brain analysis irrespective of trial condition.

## 4 Discussion

This study investigated the causal effects of disrupting the right IFG across remote nodes of the social brain network (Amodio & Frith, 2006; Frith & Frith, 2007). Participants were scanned while performing a delayed match-to-sample emotion recognition task after TMS was delivered over the right IFG or over the vertex control site. Results revealed increased activity in the bilateral anterior superior temporal sulcus (aSTS), medial prefrontal cortex (mPFC) and precuneus when TMS was delivered to the right IFG compared to the vertex (Figure 2). These findings show that the disruption of top-down control mechanisms in the lateral PFC alters the neural processing in remote nodes of the social brain network. The increase in activity in the aSTS, mPFC and precuneus indicates that the broader Theory of Mind (ToM) network compensated for the transient IFG disruption to support emotion recognition (Figure 2). The IFG contains mirror neurons which support the covert simulation of emotional states, a mechanism that has been linked to the ability for emotion recognition (Adolphs, 2002; Adolphs et al., 2000; Bastiaansen et al., 2009; Kilts et al., 2003). Covert simulation refers to the simulation of perceived facial expressions without the observer overtly mimicking the emotion (Bastiaansen et al., 2009; Jabbi & Keysers, 2008). Previous studies have found that damage to the IFG, particularly the pars opercularis, impaired simulation-based functions, such as emotional empathy (Shamay-Tsoory et al., 2009) and perceptual mentalizing, that is identifying the mental state of others based on perceptual cues such as facial expressions (Herbet et al., 2014). With a disruption to the simulation-based processing that is typically supported by the IFG, the increased activity observed in the ToM network in this study suggests that the semantic representation and conceptual knowledge of emotions are recruited as a compensatory mechanism.

Recognising emotions is not solely a perceptual process, a broader semantic representation of emotion can be seen when the concept of a particular emotion such as fear is expressed through various ways including facial expressions, words and nonverbal sounds (Carroll & Young, 2005). Emotion perception employs conceptual knowledge about emotions, this is shown in studies where seeing an emotion word primed and facilitated the perception of an emotional facial expression (Gendron et al., 2012; Lindquist & Gendron, 2013; Nook et al., 2015). This current study required maintaining and matching emotions across different types of stimuli (the word happy or fearful and a happy or fearful face), therefore delivering TMS to the right IFG before this task could have disrupted how the brain integrated the concept of an emotion that was presented across printed words and facial expressions.

Theory of mind refers to the ability to infer another person’s beliefs, motivations and emotional states. This mentalising system includes cognitive ToM, that is the ability to understand another person’s thoughts and beliefs, and affective ToM which involves the ability to infer another person’s emotional state and what they are feeling (Frith & Frith, 2006; Schlaffke et al., 2015; Schnell et al., 2011). These two types of ToM are represented in different brain regions with previous studies implicating the IFG in affective mentalising/ToM (Schlaffke et al., 2015; Schnell et al., 2011; Schurz et al., 2014). Transient disruption to the IFG before completing the emotion recognition task in this study led to compensatory recruitment of other regions in the ToM network which are involved in the conceptual representation of emotions.

The increased bilateral aSTS activation found in this study is consistent with this interpretation (Figures 2c and 2d). The response of the STS is not limited to visual perception of emotions, it also responds to voices (Belin et al., 2002; Peelen et al., 2010; Young et al., 2020). This suggests that the STS contains a broader representation of emotion as it is selective to emotions irrespective of the way in which they are presented (Peelen et al., 2010). Moreover, the anterior temporal lobes (ATL) are known to store semantic concepts and damage to the ATL results in an impaired ability to generalise conceptual knowledge across exemplars (Lambon Ralph & Patterson, 2008; Schurz et al., 2014). Relating to social cognition, the ATL also provides conceptual knowledge about social behaviours, such as “ambitious” and “loyal” (Zahn et al., 2007). Therefore, the increased aSTS activity in this study may reflect a compensatory response to support the semantic or conceptual aspects of emotion recognition when the right IFG function is disrupted.

This interpretation is further supported by increased activity in the mPFC and precuneus, two regions involved in the ToM network. Akin to the STS, the mPFC represents emotions that generalise across stimulus types, for example emotional facial expressions, body expressions and vocal expressions (Peelen et al., 2010). However, unlike the STS, the mPFC also responds to emotions that are inferred from context, even in the absence of sensory cues overtly depicting emotional stimuli, such as facial expressions, bodies and vocal intonations (Skerry & Saxe, 2014). This conceptual representation of emotion and the capacity for emotional inference facilitates the role of the mPFC in understanding and categorising the emotional and mental states of others. Alongside the mPFC, the precuneus has frequently been reported to be involved in ToM tasks (Adolphs, 2009; Saxe & Powell, 2006) and when evaluating and attributing emotions to others (Lee & Siegle, 2012; Schulte-Rüther et al., 2007). The precuneus has been implicated in perspective-taking and self-other distinction (Cavanna & Trimble, 2006; Ochsner et al., 2004), a function that is relevant to understanding the emotional state of others and acknowledging that it can differ from one’s own emotions. Together, the increased activation of aSTS, mPFC and precuneus indicates a distributed compensatory mechanism in which conceptual, evaluative, and self-other representational systems collectively support emotion recognition when the right IFG is disrupted.

This conclusion may also explain the difference in our prior study that examined the neural effects of TMS disruption to the right IFG (Pitcher et al., 2023). In this study participants silently named the expressions depicted on faces, a simpler behavioural task that required brief external focus. TMS disruption of the right IFG may have resulted in a failure of top-down suppression causing an increase of activity in the default node network as reported in a prior study that combined TMS and fMRI (Chen et al., 2013). By contrast the delayed match-to-sample task we used in the present study required both working memory and the processing of semantic information. Disruption of these processes after TMS disruption of the right IFG necessitated a greater recruitment of neural activity in nodes of the social brain (e.g. the aSTS and mPFC). The differences in the results of the two studies may then reflect a dynamic switch between passive network intrusion and active compensatory plasticity, determined by whether the concurrent task is dependent on visual perception or required more complex semantic processing.

Notably, the temporoparietal junction (TPJ), a key region of the ToM network did not show a significant difference when TMS was delivered to the right IFG compared to the vertex. This result is consistent with the cognitive demands of the task and aligns with the dual-stream model of ToM. Using the Reading the Mind in the Eyes task (RMET) to assess low-level perceptive-based mentalising and a comic strip task to assess high-level, inference-based mentalising, Herbet et al. (2014) found that low-level mentalising accuracy correlated with the degree of disconnection in the arcuate fasciculus. Specifically, the pars opercularis of the IFG was more likely to be damaged in patients with impaired accuracy in the RMET than in patients with unimpaired accuracy. The TPJ demonstrates selectivity to high-level inference-based mentalising such as reasoning about beliefs, intentions or narrative-based social scenarios (Saxe & Kanwisher, 2013). In contrast, the task in the present study required lower-level perceptual recognition of emotions rather than inferences of complex mental states. Consequently, there was no significant difference in TPJ activation across the two TMS sessions indicating the functional specificity of the perceptual and affective mentalising systems engaged by the task. Future studies could adapt our experimental design to examine the remote effects of transient disruption to the TPJ.

The results of this study provide causal evidence that regions across the ToM network compensate for transient disruption of the IFG during emotion recognition as suggested by neuropsychological studies (Dal Monte et al., 2014; Perry et al., 2017; Philippi et al., 2009). However, the absence of behavioural effects limits how the significant differences in brain activity can be interpreted. The behavioural measures in this study were insensitive to neural disruption potentially due to ceiling effects or because the task was not sufficiently sensitive to capture subtle impairments in emotion recognition or task performance. Studies have demonstrated the involvement of the IFG in cognitive functions such as working memory (Curtis & D’Esposito, 2003). Therefore, increasing the task difficulty, for example by employing a task that required active engagement of working memory mechanisms such as a one-back task (Chan & Downing, 2011; Volle et al., 2008) may elicit greater engagement of the IFG. Examining behavioural performance in tasks that include a higher level of difficulty or emotional ambiguity may help determine whether the network changes after IFG disruption also produce behavioural consequences.

In conclusion, TMS over the right IFG led to increased activation in regions of the social brain network during emotion recognition, including the anterior STS, mPFC and precuneus. These results suggest that neural systems supporting emotion recognition are sensitive to prefrontal disruption and that TMS to the right IFG evokes compensatory recruitment across distributed regions of the ToM network. This study reinforces the idea that emotion recognition is not a purely perceptual process but involves a network of regions supporting emotional understanding, perspective-taking and social inference. Overall, these findings suggest that the right IFG plays a causal role, perhaps through simulation-based mechanisms, in supporting these higher-order functions of the social brain.

## Acknowledgements

This work was supported by Leverhulme Trust Project Grant RPG-2024-389 (D.P.). D.K. is supported by the DFG (KA4683/5-1, project number 518483074) and under Germany’s Excellence Strategy (EXC 3066/1 “The Adaptive Mind”, project number 533717223).

